# Sex-specific thermoregulatory effects of estrogen signaling in *Reprimo* lineage cells

**DOI:** 10.1101/2024.12.02.626488

**Authors:** Jae W. Park, Laura R. Cortes, Norma P. Sandoval, Adriana R. Vree, Alejandra G. Baron, Kelly Vranich, Higor J. Fideles, Rosalizbeth M. Martinez, Elizabeth A. Dilday, Mia R. Hansen, Weronika Budek, Julissa I. Lopez, Laura G. Kammel, J. Edward van Veen, Stephanie M. Correa

**Affiliations:** Department of Integrative Biology and Physiology, University of California, Los Angeles, California, USA; Cypress College, Cypress, CA, USA

**Author notes:** co-first authors (Jae W. Park, Laura R. Cortes). The authors have nothing to disclose.

**Keywords:** Estrogen receptor alpha, thermogenesis, energy expenditure, *Reprimo*, metabolism, hypothalamus

## Abstract

Estrogens have considerable impact on energy homeostasis and metabolic health. In mice, signaling through estrogen receptor alpha (ERα) alters energy intake and multiple aspects of energy expenditure, effects that may be mediated by specific regions or neuronal sub-populations of the hypothalamus. This study investigates the function of ERα in neurons of the lineage that expresses *Rprm* (Reprimo), a gene we previously linked to thermoregulation in females. Here, we engineered a novel *Reprimo*^*Cre*^ mouse to selectively knock out ERα in *Rprm* lineage cells (Reprimo-specific estrogen receptor α KO; RERKO) and report changes in core temperature in female mice, with no changes in movement or food intake. RERKO females have elevated brown adipose tissue (BAT) temperature and lower tail temperature relative to controls, suggesting increased heat production and reduced heat dissipation, respectively. Developmental expression of *Rprm* was detected in the brain, but not in BAT or white adipose tissue suggesting temperature changes may be mediated by the nervous system. To confirm centrally mediated effects on temperature, we ablated *Rprm* expressing cells in the mediobasal hypothalamus and observed a reduction in core temperature relative to controls. Taken together, these results indicate that estrogen signaling in the *Rprm* lineage is critical for thermoregulation, mainly through the modulation of brown adipose tissue thermogenesis in female, but not male, mice.

## Introduction

Estrogens are potent modulators of energy homeostasis [1–3], and thus, plummeting levels of estrogen during menopause dysregulate metabolism [4]. Menopausal people experience hot flushes and reduced energy expenditure, which leads to greater adiposity, body weight, and risk of metabolic disorders [5–8]. Energy expenditure is regulated by estrogen signaling in the hypothalamus [9–13]. Recent work suggests that distinct groups of hypothalamic estrogen receptor alpha expressing (ERα+) neurons may regulate specific aspects of energy homeostasis [10,11,13]. Pinpointing these neural subsets and their precise functions can lead to a mechanistic understanding of how energy expenditure is regulated and inform the design of cell-based therapeutics tailored to specific symptoms in menopausal or obese patients.

Rodents are suitable models to study the hormonal regulation of energy expenditure. Like menopausal people, mice lacking ovaries have lower energy expenditure and increased body weight and adiposity [14], and estradiol administration blunts these changes [15]. Although ERα (*Esr1*) is found throughout the body, its expression in the central nervous system (CNS) is necessary for energy homeostasis. Mice with *Esr1* knocked out (KO) in the *Nestin-Cre* cell lineage that labels the nervous system have greater body mass, and females, in particular, eat more and show reduced energy expenditure relative to controls [9]. The neural circuitry involved in energy homeostasis includes the ventrolateral area of the ventromedial nucleus (VMHvl), the medial preoptic area (MPO), and the arcuate nucleus (ARH). These regions contain abundant expression of ERα. Female mice and rats with *Esr1* knockdown in the VMHvl have lower activity levels, basal metabolic rate, and energy expenditure, resulting in higher body weight relative to controls [16,17]. Similarly, mice with *Esr1* KO in the steroidogenic factor 1 (*Sf1/Nr5a1)*-lineage have lower metabolic rates and thermogenesis [9]. In contrast, activating *Esr1*+ neurons in the VMHvl (VMHvl^*Esr1*^) increases heat production and movement [11]. Thus, the VMHvl is essential for the regulation of energy expenditure. The effects of estrogens on food intake are regulated by other regions, including the nearby ARH. Knocking out *Esr1* in pro-opiomelanocortin (POMC) lineage cells increases food intake and body weight in females relative to controls [9]. Similarly, silencing steroid and metabolic hormone-responsive ERα/Kisspeptin+ neurons in the ARH causes obesity in female mice and alters circadian feeding patterns [18–21]. Lastly, activation of ERα+ cells in the MPO leads to a hypometabolic state akin to torpor; temperature, energy expenditure, metabolic rate, and movement plummet [12]. In summary, ERα+ cells or expression in the VMHvl and ARH promote energy expenditure, while *Esr1* expression in ARH cells reduce energy intake, and ERα+ MPO cells trigger a profound energy conservation state.

Intermingled within the same nuclei, ERα+ cells may differ in their function, connections, and neurochemical identity (*i*.*e*., their gene expression pattern) [22,23,11,24]. For example, a study using preoptic area slices from rats showed that a portion of estrogen-responsive MPO neurons (likely expressing ERα) are temperature-responsive, with some responding to warmth and others to cold [25]. In a similar fashion, VMHvl^*Esr1*^ cells differ in their response to glucose and their neural connectivity [26,24]. About half are excited by an increase in glucose levels and project to the dorsal raphe nucleus, whereas the other half are excited by a depletion of glucose levels and project to the arcuate nucleus [24]. Functional specialization among ERα+ cells suggests that they may differ in the expression of peptides and proteins that shape their roles and responses. Indeed, our lab found that ERα+ VMHvl cells are neurochemically heterogenous. Using single-cell sequencing, we found sub-populations of ERα+ VMHvl neurons marked by the expression of substance P (Tachykinin 1; *Tac1*), reprimo (*Rprm*), or prodynorphin (*Pdyn*). Intriguingly, females have many more ERα neurons that express *Tac1* and *Rprm* than males, and manipulating these cells differentially alters energy expenditure. Mice that are missing a subset of ERα+ VMHvl cells that co-express *Tac1* and *Nkx2-1* or have inactivation of the melanocortin receptor 4 (*Mc4r*) subpopulation move less [10,13]. With *Tac1/Nkx2-1* cell loss, this also leads to increased adiposity and body weight [10]. Interestingly, *Rprm* knockdown in the VMHvl leaves movement intact, but increases core temperature and thermogenesis. These findings suggest specialization of cell populations or gene function within estrogen sensitive hypothalamic regions.

The function of *Rprm*, a tumor suppressor gene, and *Rprm*+ neurons has been largely unexplored outside of pathological contexts [27–29], with the exception of one developmental gene expression study [30]. In the present work, we test the function of *Rprm+* cells on energy expenditure. Specifically, we hypothesize that *Esr1* expression in *Rprm+* cells modulates body temperature via thermogenic brown adipose tissue (BAT) [31]. We generated a knock-in mouse that expresses Cre recombinase under the control of *Rprm* to selectively knock out *Esr1* in *Rprm* lineage cells. Despite the extensive effects of global and CNS-specific *Esr1* knockout on weight, food intake and movement [9,32,33], *Esr1* KO in *Rprm* lineage cells produces a more restricted phenotype. Core temperature is altered between controls and KO, while body weight, body composition, and food intake are largely unaffected. Furthermore, BAT mass is heavier and produces more heat, indicating increased thermogenesis. Because *Esr1* and *Rprm* are co-expressed in several brain areas and peripheral organs, we next determined the specific role of *Rprm* cells in the mediobasal hypothalamus (MBH) using a Cre-dependent caspase. Loss of *Rprm* cells led to lower core temperature and higher BAT mass compared with controls. In summary, our data suggest that *Esr1* expression in *Rprm* lineage cells and MBH^*Rprm*^ neurons seem to largely modulate core temperature, while leaving other aspects of energy balance intact.

## Methods

### Generation and validation of a Rprm^Cre^ knock-in mouse

The donor construct consisted of a double stranded DNA repair cassette that included the coding sequence for *Rprm* linked to the codon-improved cre recombinase (*iCre*) sequence via a P2A peptide (Fig. 1A). The P2A peptide leads to ribosomal skipping and the generation of distinct peptide products, in roughly equal amounts, from a single multi-cistronic construct [34]. A 21-bp nuclear localization sequence (NLS) is located at the 5’ end of the *iCre* exon and helps direct proteins into the nucleus. To induce proper DNA insertion, we also included homology arms (approximately 2,000 bp) both upstream and downstream of the *Rprm* and *iCre* coding sequences. Finally, 34-bp long flippase recognition target (FRT) sites were inserted in the upstream and downstream homology arms within 250 bp of the 5’ and 3’ exons. The total length of the construct is approximately 8.3 Kb. The engineered DNA sequences were based on the C57BL/6J mm8 reference genome and introduced into albino C57BL/6 blastocysts by CRISPR injection by the University of California San Diego Transgenic & CRISPR Mouse Core. Two founder lines were generated but only one was confirmed to contain the entire donor sequence by PCR validation and sequencing. The founder was crossed to C57BL/6J females and heterozygous offspring were used to generate a homozygous line using in-crosses.

**Figure 1:**
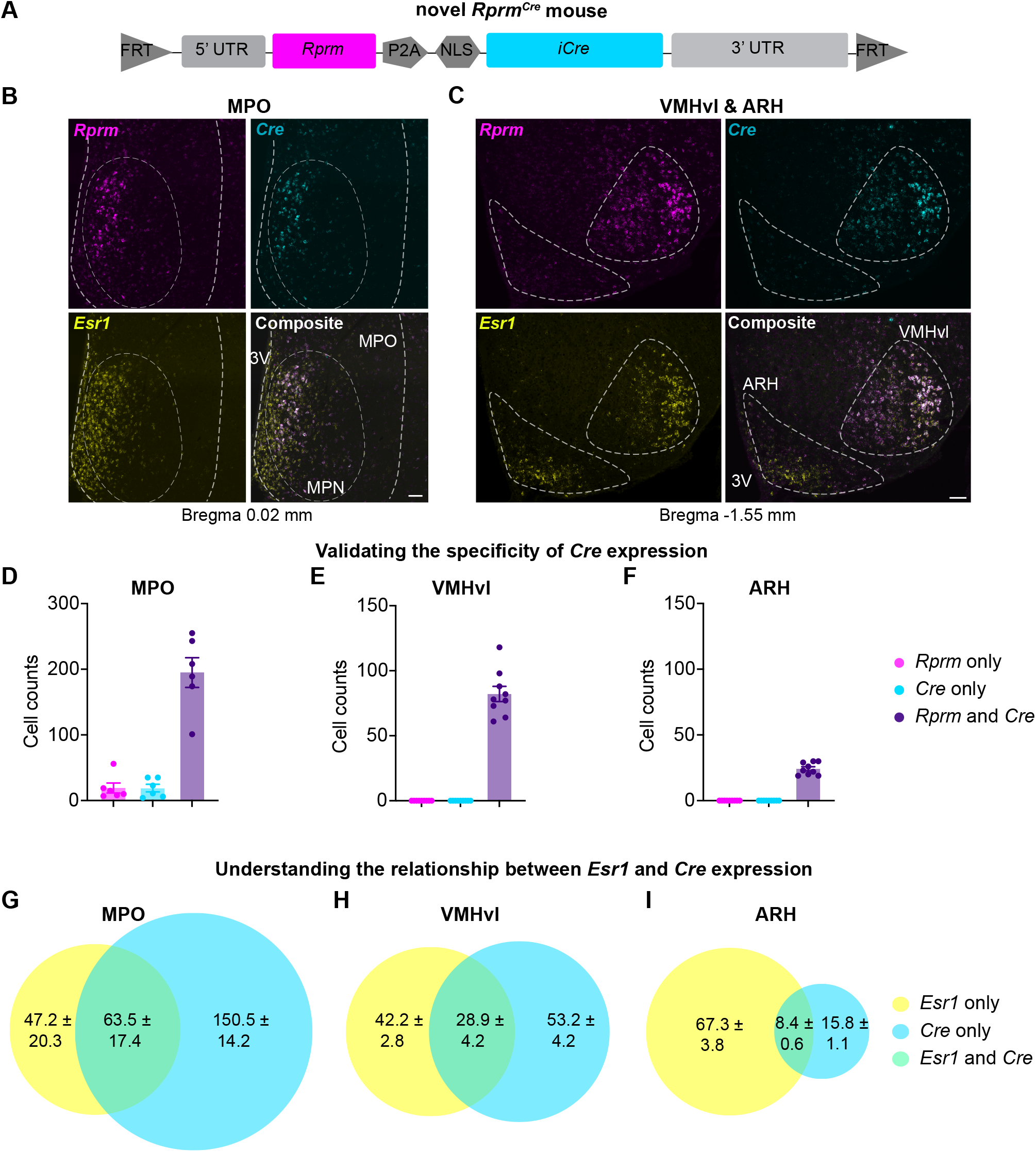
Characterization of a novel *Rprm*^*Cre*^ knock-in mouse model. (A) Schematic of the knock-in construct. FRT = flippase recognition target sites, UTR = untranslated region, NLS = nuclear localization sequence. (B, C) Single-molecule *in situ* hybridization labeling *Rprm* (magenta), *Cre* (Cyan) and *Esr1*^*+*^ (yellow) cells in coronal sections of the brain containing the (B) MPO and (C) VMHvl & ARH in female mice. (D-F) Quantification of cells expressing *Rprm* alone (magenta), *Cre* alone (*Cyan*), and both *Rprm* and *Cre* (purple) in the MPO (D), VMHvl (E), and ARH (F) show that more than 91% of *Rprm*^*+*^ cells express *Cre* (n = 6-9 per region). (G-I) Average number of cells expressing *Esr1* alone (yellow), *Cre* alone (Cyan), and both *Esr1* and *Cre* (green) are depicted in the MPO (G), VMHvl (H), and ARH (I; n = 6-9 per group). The mean ± SEM are depicted in D-I, as well as individual data points in D-F. Scale bar = 100 μm. MPO = medial preoptic area, MPN = medial preoptic nucleus, VMHvl = ventrolateral area of the ventromedial nucleus of the hypothalamus, ARH = arcuate nucleus, 3V = third ventricle.

*Esr1* ^f/f^ female mice [35] were paired with *Rprm*^*Cre*^;*Esr1*^f/f^ studs to generate Reprimo-specific Estrogen Receptor α Knock **<u>O</u>**ut (RERKO) mice on a mixed B6;129P background. In this RERKO line, mice had *Esr1* knocked out (KO) in *Rprm* lineage cells. Mice were bred and maintained in approved conditions at the University of California Los Angeles. Control littermates were *Esr1*^f/f^ and denoted WT for the *Rprm*^*Cre*^ mutation. For lineage experiments, *Rprm*^*Cre*^ mice on a C57BL/6J background were paired with *Ai14* ^f/f^ reporter mice (Jax #007914; [36]) to generate offspring that fluoresce tdTomato in tissues expressing *Rprm* (*Rprm*^*Cre*^; *Ai14* ^f/+^). Mice were kept in 12:12 light cycle conditions (zeitgeber time (ZT) 0 = 7 am) at ∼23 °C and ∼30% humidity with food and water provided *ad libitum*. All animal studies were approved by the UCLA Institutional Animal Care and Use Committee (IACUC) and were carried out following the recommendations of the Guide for the Care and Use of Laboratory Animals of the National Institutes of Health.

### Surgeries

For all surgeries, mice were anesthetized with isoflurane and administered buprenorphine (0.01 mg/mL) and carprofen (0.58 mg/mL) on the day of surgery and the following day to manage pain and inflammation.

### Probe implants

G2 eMitters (Starr Life Sciences; Oakmont, PA, USA) were placed into the abdominal cavity to record core temperature and locomotion. Cubisens TS100/TS110 probes (Cubeworks; Ann Arbor, MI, USA) or Nano-T loggers (Star-Oddi, Garðabær, Iceland) were secured subcutaneously above the BAT depots or over the base of the tail to gauge heat generation and dissipation, respectively. Custom sleeves were designed to secure probes above BAT and tail using Autodesk Fusion 360 (San Francisco, CA, USA) and UltiMakerCura software (New York, NY, USA). Sleeves were printed on a Creality Ender-2 V2 3D printer using PLA filament. Files are publicly available on NIH 3D (Reference model: 3DPX-020322; https://3d.nih.gov/entries/3DPX-020322).

### Stereotaxic surgery

Control (*pAAV8-Syn-FLEX-Mac-GFP*) or caspase virus (*pAAV2-flex-taCasp3-TEVp*) was delivered into the VMHvl (AP 1.58, ML ± 0.65, DV 5.7) in 8-week-old C57BL/6J wildtype or *Rprm*^*Cre*^ mice [37,38]. Each hemisphere received 100 nL at 5-10 nL/sec using a 10 μL Hamilton syringe attached to a pulled glass needle. Mice recovered for two weeks before starting three days of temperature and locomotion recordings which were averaged hourly. Targeting of the VMHvl was validated by immunohistochemical labeling of ERα (described below), since there is no suitable reprimo antibody and brains were not collected RNAse-free. Because it could not be confirmed that *Rprm*+ cells were unaffected in nearby nuclei, such as the ARH, this manipulation is termed MBH-specific.

### Collection

Mice were perfused with 4% paraformaldehyde (PFA; Electron Microscopy Sciences; Hatfield, PA, USA) in 0.01M phosphate-buffered saline (1x PBS). Brains were stored in 4% PFA overnight at 4°C and transferred to 30% sucrose in 1x PBS until brains sank. Brains were embedded in Optimal Cutting Temperature compound (OCT; Fisher Scientific, Fair Lawn, NJ, USA) and stored at −80°C until cutting. BAT, inguinal white adipose tissue (iWAT), and gonadal white adipose tissue (gWAT) were dissected and stored in tissue cassettes submerged in 70% ethanol until staining.

### Metabolic and reproductive phenotyping

Mice were group housed according to sex, and body weight was recorded weekly from 5 to 16 weeks of age. A separate cohort of mice were weighed and placed into an EchoMRI™ (Houston, TX, USA) to record fat and lean mass at 16 weeks of age. To record food intake, a pre-weighed allotment was provided to singly housed mice daily for 7 consecutive days. Every 24 hours, the remaining food and any crumbs in the cage were weighed and subtracted from the amount initially provided to determine daily food intake, which was averaged over seven days to provide a daily average of food consumption. To record core temperature and locomotion, mice were placed on ER4000 energizer/receiver pads in their home cages (version 5, VitalView software, Starr Life Sciences). All temperature and activity measurements were collected every five minutes across three to five days. For temperature and activity recordings, an average for each hour of the day (ZT 0 – 23) across recording days was calculated. Raw data from G2 emitters, TS100/TS110 probes, and Nano-T loggers were exported in CSV format using VitalView (Starr Life Science), TS GUI (Cubeworks), and Mercury (Star-Oddi), respectively. Data were prepared for analysis in Microsoft Excel (Microsoft Corporation; Redmond, Washington, USA). R Studio (R version 4.3.1; Boston, MA, USA) was used to process the data and generate hourly averages (code for processing and plotting provided at https://github.com/kodori00/Rscript). To track the estrous cycle, mice underwent vaginal lavage using 0.9% saline every morning between ZT 1-2.

### Histology

#### Cytology

For estrous cycling, vaginal lavages were deposited onto slides, stained with Giemsa (0.6% in 1x PBS;) to visualize nucleic acid and distinguish cell morphology, and imaged using light microscopy as described in Massa et al., 2023. The estrous stage was determined by the relative amounts of leukocytes and cornified and nucleated epithelial [39]. BAT and iWAT were sectioned into 4 μm slices and stained with hematoxylin and eosin by the UCLA Translational Pathology Core Laboratory.

#### Immunohistochemistry

Brains were coronally cryo-sectioned into four or six 20-30 μm series and stored on slides or in cryoprotectant (30% sucrose, 30% ethylene glycol, and 1% polyvinylpyrrolidone in 1x PBS) at −80°C until use. One series of 30 μm thick sections underwent immunohistochemistry to validate the loss of ERα expression in RERKO mice (rabbit anti-ERα, 1:1000, ThermoFisher Scientific, Waltham, MA; Cat# PA1-309, RRID:AB_325814) and one series of 20 μm sections was used to validate the loss of ERα cells in caspase-treated mice (rabbit anti-ERα, 1:500, Millipore Sigma, Burlington, MA; Cat# 06-935, RRID:AB_310305). Tissue was rinsed in 1x PBS and blocked for one hour at room temperature (RT) in either 10% bovine serum albumin (BSA) and 2% Normal Goat Serum (NGS) or 10% NGS in 0.3% PBS-Triton X depending on antibody. After blocking, tissue was incubated in primary antibody overnight at 4°C or RT. The following day, the tissue was incubated with either Alexa Fluor 546 (1:500; Thermo Fisher Scientific Cat# A-11035, RRID:AB_2534093) or 647 (1:1000; Jackson ImmunoResearch Labs, West Grove, PA; Cat# 111-605-144, RRID:AB_2338078) anti-rabbit secondary antibody for 1.5-2 hours at RT. Nuclei were counterstained with 4′,6-diamidino-2-phenylindole (DAPI, 1:500-1000, ThermoFisher Invitrogen), and Fluoromount-G was used to coverslip (Southernbiotech, Birmingham, AL, USA).

#### Fluorescent In-Situ Hybridization

Perfused brains (4% PFA) from female mice were coronally cryo-sectioned into six series at 20 μm. Sections containing the MPO, VMH, and ARH were anatomically-matched across mice.

Fixed tissue underwent single molecule *in-situ* hybridization according to manufacturer instructions in the RNAscope Multiplex Fluorescent Detection Kit version 2 (Advanced Cell Diagnostics, Newark, CA, USA). Tissue was fixed in 10% neutral buffered formalin for 20 minutes and pre-treated using serial ethanol dilutions, hydrogen peroxide, and protease III solution (incubation time shortened to 17 minutes to prevent tissue degradation). Tissue was incubated with probes targeting *Rprm* (Cat# 466071), *Cre* (Cat# 1058921), and *Esr1* (Cat # 478201) at 40°C for 2 hours. Afterward, the probe signal was amplified and fluorescently labeled with Opal 520 (1:750), 570 (1:500), and 690 (1:500; Akoya Biosciences, MA) for *Esr1, Rprm* and *Cre*, respectively. Signal was developed using horseradish peroxidase and counterstained with reagents provided in the kit. Slides were cover-slipped using Prolong Gold Antifade (ThermoFisher Scientific).

#### Imaging & analysis

Brightfield images of BAT and iWAT histological sections were taken by a Leica DM1000 microscope (Leica Microsystems Inc, Deerfield, IL, USA). Brain, uterine and ovarian sections were imaged using a Nikon Eclipse Ti2 inverted microscope. Peripheral tissues were illuminated using a Leica MZ10F and imaged using a dual MP camera system on an iPhone13. ERα cells were counted in the anteroventral periventricular area (AVPV), medial preoptic area (MPO), ventromedial ventrolateral area (VMHvl), and arcuate nucleus (ARH) of the hypothalamus. The machine learning software Ilastik (Berg et al., 2019; www.ilastik.org/about) segmented ERα+ cells from the background. Positive object outlines were loaded to CellProfiler 4.2.5 (Stirling et al., 2021; www.cellprofiler.org), overlaid over the fluorescent images, manually edited if necessary, and counted. Cell counts were averaged across hemispheres; if one hemisphere was damaged, the count on the intact side was used.

For *in situ* hybridization, cell counts in the MPO, VMH, and ARH were done in one hemisphere using ImageJ Version 2.1.0/1.53c. A cell was considered positive for a given gene if it contained more than approximately 7 puncta and co-localized with DAPI. For studies interrogating the co-localization of *Cre*/*Rprm* or *Cre*/*Esr1*, each channel was analyzed independently and channels were then merged to identify co-expressing cells. For all analyses, regions of interest were determined by examining the shape of the anterior commissure, median eminence, and third ventricle referencing the Allen Brain Mouse (v2, 2011) and Paxinos and Franklin (v3, 2008) atlases. Area-proportional Venn diagrams, generated using BioVenn [40], show the mean number of cells expressing *Esr1* only, *Cre* only, or *Esr1* and *Cre* in each brain region. To estimate the proportion of the *Esr1*+ population that expresses *Rprm-Cre*, the number of *Esr1*+/*Cre*+ cells was divided by the total number of *Esr1*+ cells (*Esr1*+/*Cre*+ and *Esr1*+/*Cre*-) and multiplied by 100 to get a percentage. This proportion was calculated per animal, averaged across the group, and described in text as an average ± SEM.

#### Statistics

Statistical analyses and plots were made in Prism (version 10.2.3 for Mac; GraphPad Software, Boston, MA, USA). When applicable, two-way ANOVAs or mixed-effects models were run with repeated (age or ZT) and independent (genotype or sex) factors. Post hoc comparisons between control and RERKO mice at each time point were performed using Šídák’s multiple comparisons test. Comparisons between two samples were tested with t-tests, as indicated in the figure legends.

## Results

### 3.1 Validation of *Rprm*^*Cre*^ mouse

We bred *Rprm*^*Cre*^ mice with *Ai14* reporter mice to identify the tissues expressing *Rprm* at any point through development. We report *Cre*-mediated expression of TdTomato in the brain, uterus, ovary, pancreas and pituitary (Supp. Fig. 1A,B). Conversely, TdTomato expression was absent from BAT, iWAT, and gWAT (Supp. Fig. 1C). We note that the brain exhibited widespread TdTomato expression that is most striking in the cortex. This was somewhat expected as *Rprm* expression is pronounced in the cortex and extends to many regions in development. Notably, *Rprm* expression has an interesting developmental pattern, peaking just before birth (E18.5) and dampening down by weaning (Allen Institute for Brain Science developing mouse brain, ISH data, entrez ID: 67874).

Therefore, to validate the fidelity of the transgenic allele, we assessed the expression of *Cre* and *Rprm* transcripts in adult mice. We analyzed three regions that abundantly express *Esr1*: the MPO, VMHvl, and ARH (Fig. 1B-F). In the VMHvl and ARH, *Cre* expression was exclusively restricted to *Rprm*+ cells (Fig. 1E,F). This was also the case in the MPO, except for a small number of *Cre+/Rprm-* and *Rprm+/Cre-* cells (Fig. 1D). Next, we quantified the number of *Esr1*+, *Cre*+ and co-expressing *Esr1*+/*Cre*+ cells (Fig. 1G-I) and calculated the proportion of *Esr1*+ cells that co-express *Cre*+ (and by extension, *Rprm*), reporting 58.9 ± 6.0 % in the MPO, 39.5 ± 3.4 % in the VMHvl, and 11.3 ± 0.9 % in the ARH, consistent with previous work [11]. Finally, we confirmed that the *Cre* knock-in allele does not exhibit ectopic *Rprm* expression in the MPO and VMHvl by finding no difference in *Rprm*+ cell counts between *Rprm*^*Cre*^ and WT C57BL/6J controls (Supp. Fig 2). Together, these results validate the fidelity of the expression of the *Rprm*^*Cre*^ allele in adults. Therefore, the widespread expression observed with lineage tracing likely reflects transient expression of *Rprm* throughout the brain and other tissues that extinguishes prior to adulthood.

### 3.2 *Esr1* knockout in *Rprm* lineage cells (RERKO) leads to an increase in brown adipose 0ssue mass, specifically in females

Next, we knocked out *Esr1* in *Rprm*-lineage cells which, based on TdTomato and expected *Esr1* expression, is expected to occur in the hypothalamus, uterus, ovary, pancreas, and pituitary [41,42]. While previous studies report that global ERα knockout or VMHvl-specific ERα knock-down increases body weight, feeding, and fat mass in mice [16,32], we find that body weight in RERKO mice is largely unchanged, relative to controls (Supp. Fig. 3). Male RERKO mice became subtly heavier as they aged (genotype x age interaction: F_11,318_ = 2.398, *p* < 0.0001), while females did not differ from controls. However, we observed a sex-specific effect of RERKO on BAT mass (genotype x sex interaction: F_1,40_ = 24.5, *p* < 0.0001; Fig. 2A) and no effect on iWAT (Fig. 2B). Female RERKO mice had larger BAT depots than controls (*p* < 0.0001), while BAT size in males did not differ (Fig. 2A). BAT tissue in RERKO mice appeared lighter in color and typically had larger adipose droplets relative to controls (Fig. 2C, D) – although an analysis comparing nuclei and adiposity in BAT did not reach significance (Supp. Fig. 4). In contrast to global *Esr1* KO [16,32], RERKO mice have no changes in adiposity: body mass and percent fat and lean mass as recorded via EchoMRI were similar between RERKO and control littermates in a separate cohort of animals (Fig. 2E-G). In addition, RERKO mice did not consume more food relative to controls (Fig. 2H) as reported in CNS-restricted *Esr1* KO mice [9]. Together, these data suggest that *Esr1* in *Rprm* lineage cells may regulate discrete facets of metabolism (*i*.*e*., BAT), while sparing energy intake and storage.

**Figure 2.**
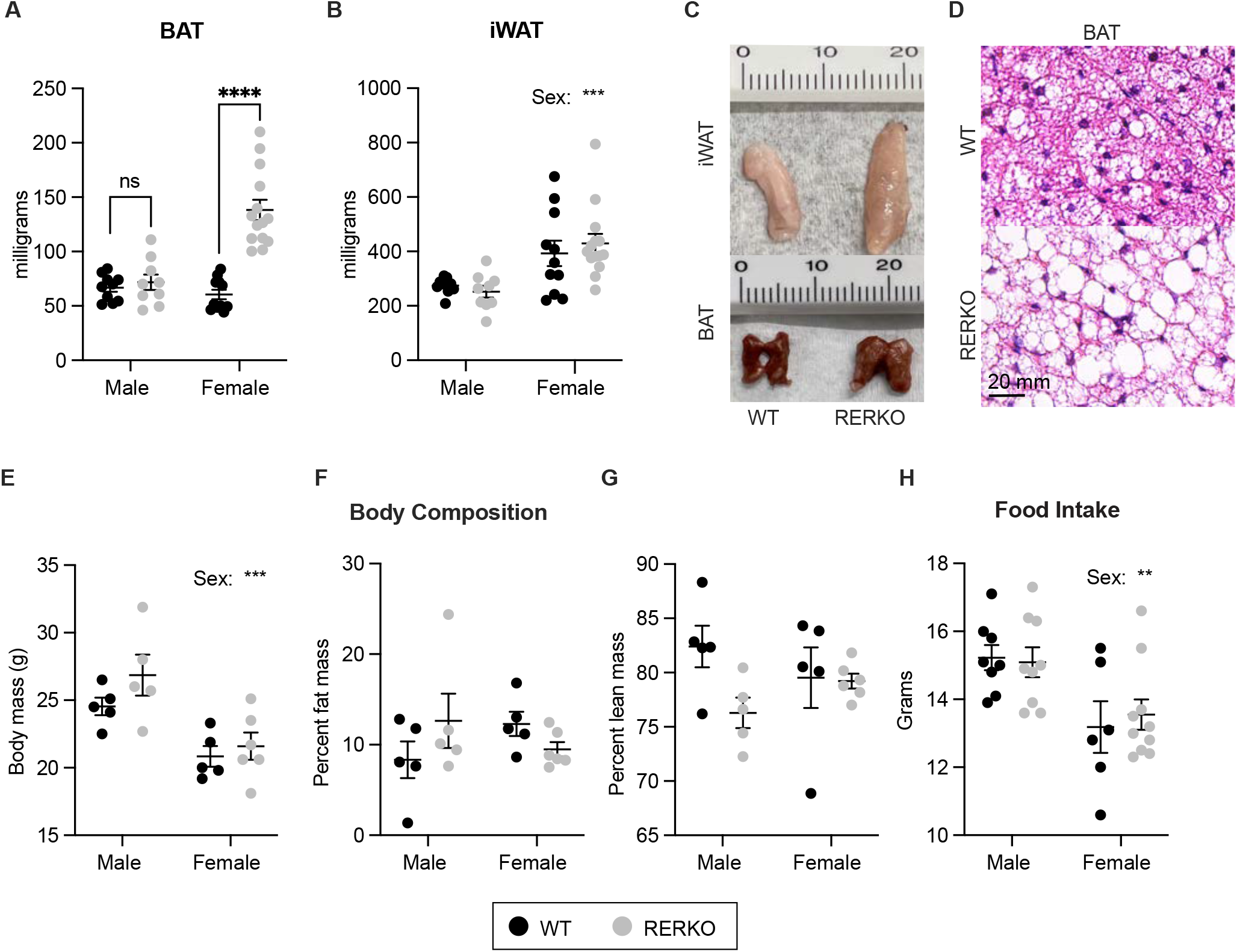
*Esr1* KO in *Rprm* lineage cells (RERKO) sex-specifically alters brown adipose tissue. (A,B) BAT (A) and iWAT (B) mass from adult male and female control littermates (black dots) and RERKO mice (gray dots). N = 10 WT males, 11 WT females, 9 RERKO males, and 14 RERKO females. (C) Representative images of iWAT (top) and BAT (bottom) in WT (left) and RERKO (right) female mice. Ruler with centimeters added for scale. (D) Representative images of hematoxylin and eosin staining of WT (top) and RERKO (bottom) in 8-week-old mice. Scale = 20 μm. (E-G) Body composition analysis of male and female WT and RERKO mice at 16 weeks of age via EchoMRI: body mass (E), percent fat mass (F), and percent lean mass (G). N = 5 WT males, 5 WT females, 5 RERKO males, and 6 RERKO females. H) Daily food consumption, accounting for spillage, averaged over seven days in WT and RERKO mice. N = 8 WT males, 6 WT females, 9 RERKO males, and 10 RERKO females. The mean ± SEM are depicted on all graphs, as well as individual data points. Two-way ANOVAs used for statistical analysis; differences between WT and RERKO tested within each sex using Šídák’s multiple comparisons test following a significant interaction. ** *p* < .01, *** *p* < 0.001, **** *p* < 0.0001.

### 3.3 *Esr1* knockout in *Rprm* lineage cells sex-specifically impacts energy expenditure

Because BAT contributes to thermoregulation, we next tested if RERKO mice had changes in body temperature by implanting telemetry probes that simultaneously and passively record core, BAT, and tail temperature (Fig. 3A). In males, core temperature and locomotion were unaffected by genotype (Fig. 3B, C), consistent with no change in BAT mass. In contrast, both temperature (genotype x ZT: F_23,437_ = 6.0, *p* < 0.0001; Fig 3D) and locomotion (ZT x genotype: F_23,391_ = 1.7, *p* = 0.02; Fig. 3E) differed between WT and RERKO female mice. Probing further, we compared BAT and tail temperature as proxies for heat generation and dissipation, respectively. Female RERKO mice had higher BAT temperature than WT mice (main effect of genotype: F_1,14_ = 4.7, *p* < 0.05; Fig. 3F) and lower tail temperature than WT mice (main effect of genotype: F_1,6_ = 8.6, *p* < 0.05; Fig. 3G). Hence, female RERKO mice appear to produce more heat and dissipate less heat, contributing to their altered core temperature phenotype. In line with the morphological findings, ERα in *Rprm+* cells appears to specifically regulate thermogenesis in a sex-specific manner.

**Figure 3.**
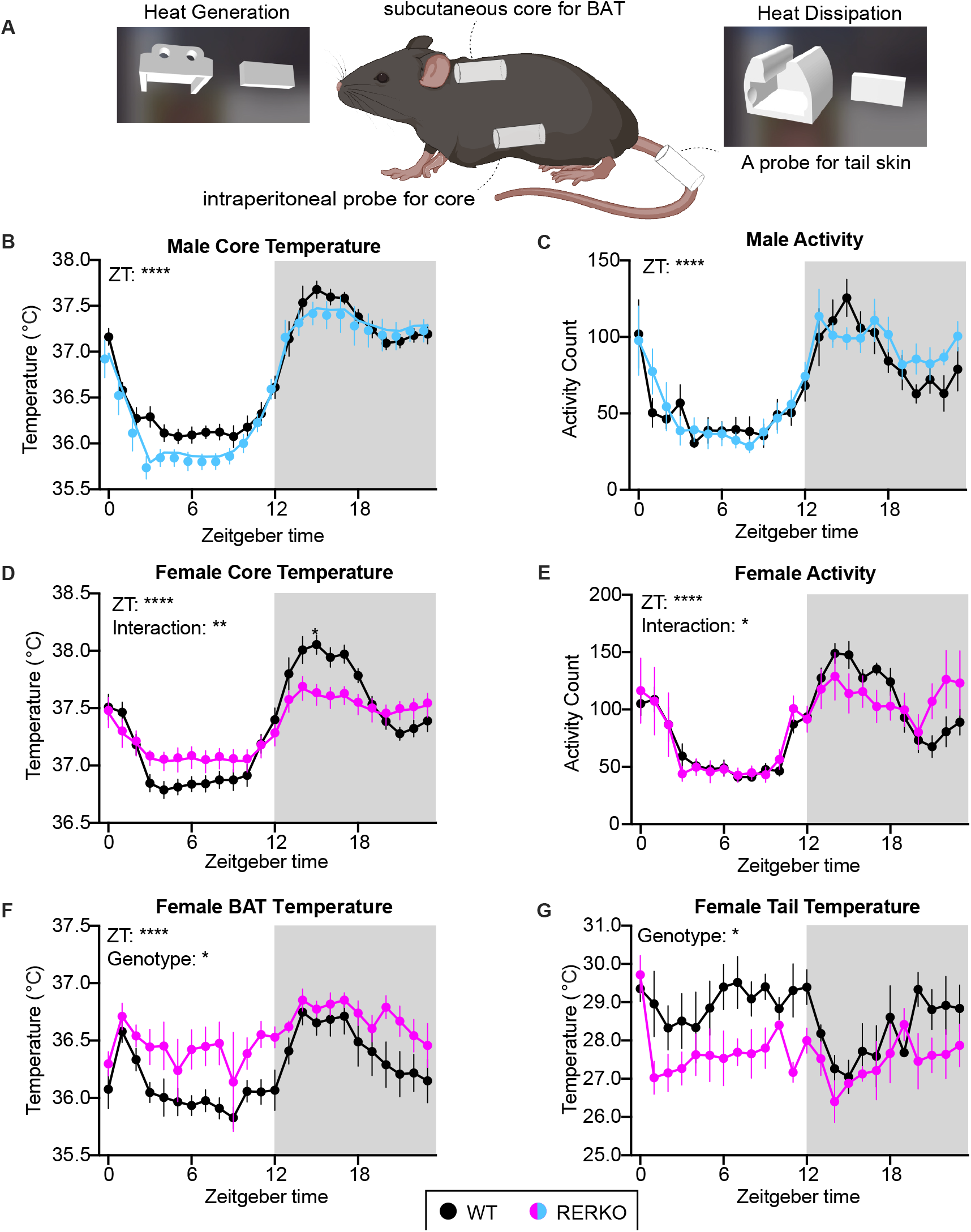
KO of *Esr1* in *Rprm* lineage cells sex-specifically impacts thermoregulation. (A) Illustration of a mouse implanted with an intraperitoneal probe to record temperature and locomotion. In addition, a probe is placed above the BAT and on the base of the tail to measure heat generation and heat dissipation, respectively. Custom 3d printed sleeves are used to secure the probes. (B-E) Temperature in Celsius or activity counts of WT siblings (black) and RERKO mice (males: blue, females: pink) averaged hourly across 24 hours. N = 8 WT males, 8 RERKO males, 11 WT females, 10 RERKO females. (F, G) Temperature averaged hourly above the BAT or at base of the tail in female control (black) and RERKO (pink) across 24 hours. N for BAT = 8 WT, 8 RERKO; N for tail = 4 WT, 4 RERKO. Mean ± SEM plotted at each time point. Two-way mixed ANOVAs used for statistical analyses. Significant interaction followed by Šídák’s multiple comparisons at each time point. ZT = Zeitgeber time. C = Celsius. Shaded box indicates the dark (active) phase of the day.

In addition to altered thermogenesis, RERKO female mice appeared to have reproductive deficits. RERKO mice did not cycle well, mostly remaining in diestrus or metestrus (Supp. Fig. 4A, B). RERKO mice had heavier ovarian (t_8_ = 2.5, p < 0.05) and decreased uterine masses (t_23_ = 5.9, p < 0.0001; Supp. Fig. 5D), and histological examination revealed abnormal ovarian and uterine anatomy (Supp. Fig. 5C), including the presence of follicular cysts and the absence of corpora lutea. Sex steroid hormones influence thermoregulation [43,44]; thus, it is possible that differing hormonal milieu between WT and RERKO underlie the temperature phenotype. To address this, females were ovariectomized, and temperature was compared between WT littermates and RERKO mice seven days later. There was a main effect of genotype (F_1,12_ = 5.5, *p* < 0.05) and an interaction between genotype and hour of day (F_23,270_ = 1.7, *p* < 0.05; Supp. Fig. 6). Ovariectomized RERKO mice had elevated temperature relative to ovariectomized controls, particularly during the light phase (Supp. Fig. 6). Thus, RERKO mice have altered thermoregulation, that is at least partially, independent from changes in ovarian hormones.

### 3.4 *Esr1* knockout in *Rprm* cells reduces ERα immunoreactivity in thermoregulatory hypothalamic regions

In the brain, ERα is expressed in brain regions involved in thermoregulation and energy intake and expenditure (*e*.*g*., MPO, VMHvl, and the ARH) [12,45], and *Rprm* is co-expressed in all of these regions (Fig. 1). RERKO mice had fewer ERα-expressing cells in the AVPV, VMHvl, and ARH (Fig. 4D, F, G), but not in the MPO (Fig. 4E). We note that the sex difference in ERα in the VMHvl was completely abolished in RERKO mice, due to a reduction in both sexes (Figure 4F). We did not have sufficient power to confirm the expected sex difference in ERα cell counts in the AVPV, but nevertheless, observed a decrease in RERKO mice relative to controls in males and females (Fig. 4D).

**Figure 4.**
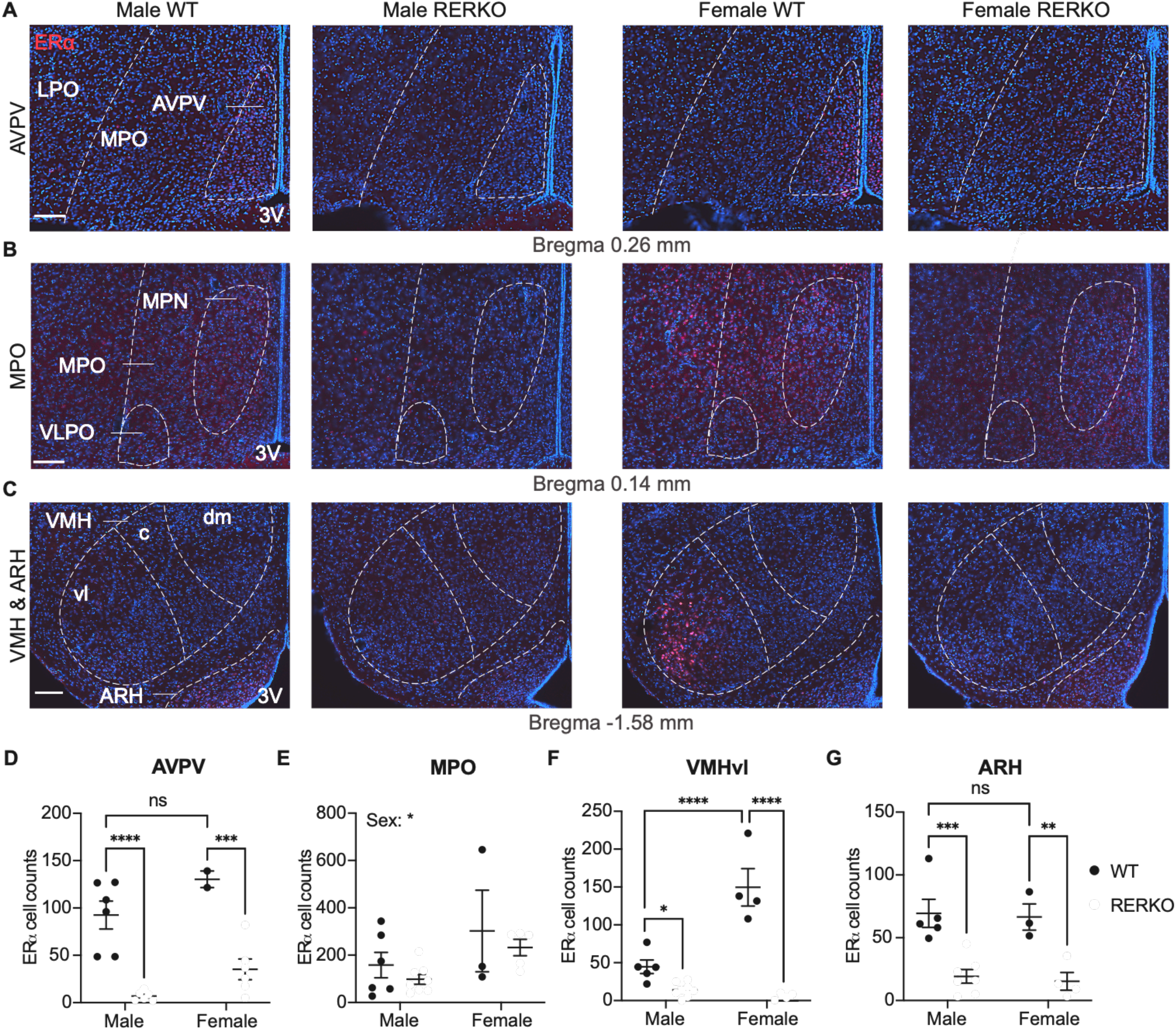
RERKO mice have reduced ERα immunoreactivity in hypothalamic thermoregulatory hubs. (A-C) Images of ERα immunoreactivity (red) in coronal sections of the brain containing the anteroventral periventricular area (A; AVPV), medial preoptic area (B; MPO), ventrolateral area of the ventromedial hypothalamus (C; VMHvl), and arcuate nucleus (C; ARH). Tissue is counterstained with DAPI (blue). (D-G) Quantification of ERα+ cells in male and female WT (black dots) and RERKO (gray dots) mice in the AVPV (D), MPO (E), VMHvl (F) and ARH (G) (N = 2 - 8 per group). Statistical analysis performed using Two-way ANOVA followed by Šídák’s multiple comparisons test, if applicable. The mean ± SEM are depicted. Individual data points are shown in figures. ** *p* < .01, *** *p* < 0.001, **** *p* < 0.0001. Scale bars = 100 μm. Boundaries of nuclei and landmarks outlined to aid in visualization. LPO = lateral preoptic area, MPN = medial preoptic nucleus, VLPO = ventrolateral preoptic area. dm = dorsomedial, c = caudal, 3V = third ventricle.

### 3.5 Ablating *Rprm*+ cells in the MBH decreases core temperature and increases fat depots

We hypothesized that the temperature phenotype in RERKO mice may involve the VMHvl given our previous work [11]. To test this, we delivered an AAV expressing a Cre-dependent and self-activating caspase into the VMHvl of *Rprm*^*Cre*^ mice to ablate *Rprm+* cells in this brain region [38]. We labeled ERα to validate caspase-mediated cell death of *Esr1*/*Rprm* cells in the VMHvl and ARH and observed decrease in ERα+ immunoreactivity between controls and caspase-treated mice in the VMHvl, but not ARH (Figure 5C). Although we did not detect a decrease in ERα+ cell counts in the ARH, we note that this does not rule out the possible loss of other *Rprm*+ cells in a broader area of the MBH. Caspase-treated mice had decreased core temperature (main effect of group: F_1,10_ = 11.20, *p* < 0.01; Figure 5D) and heavier BAT (t_10_ = 4.02, *p* < 0.01, Figure 5G), iWAT (t_10_ = 2.23, *p* = 0.05; Figure 5H) and gWAT (t_12_ = 4.03, *p* < 0.01; Figure 5I). Statistical differences in activity levels (Figure 5E), total body mass (Figure 5F), or uterine mass (Figure 5J) were not detected. This provides additional evidence that *Rprm*+ neurons in the MBH regulate body temperature [11]. Importantly, we observe these effects without an effect on the uterus, a bioassay for circulating estrogen levels [46], suggesting that this effect is not secondary to any potential changes in circulating estrogens.

**Figure 5.**
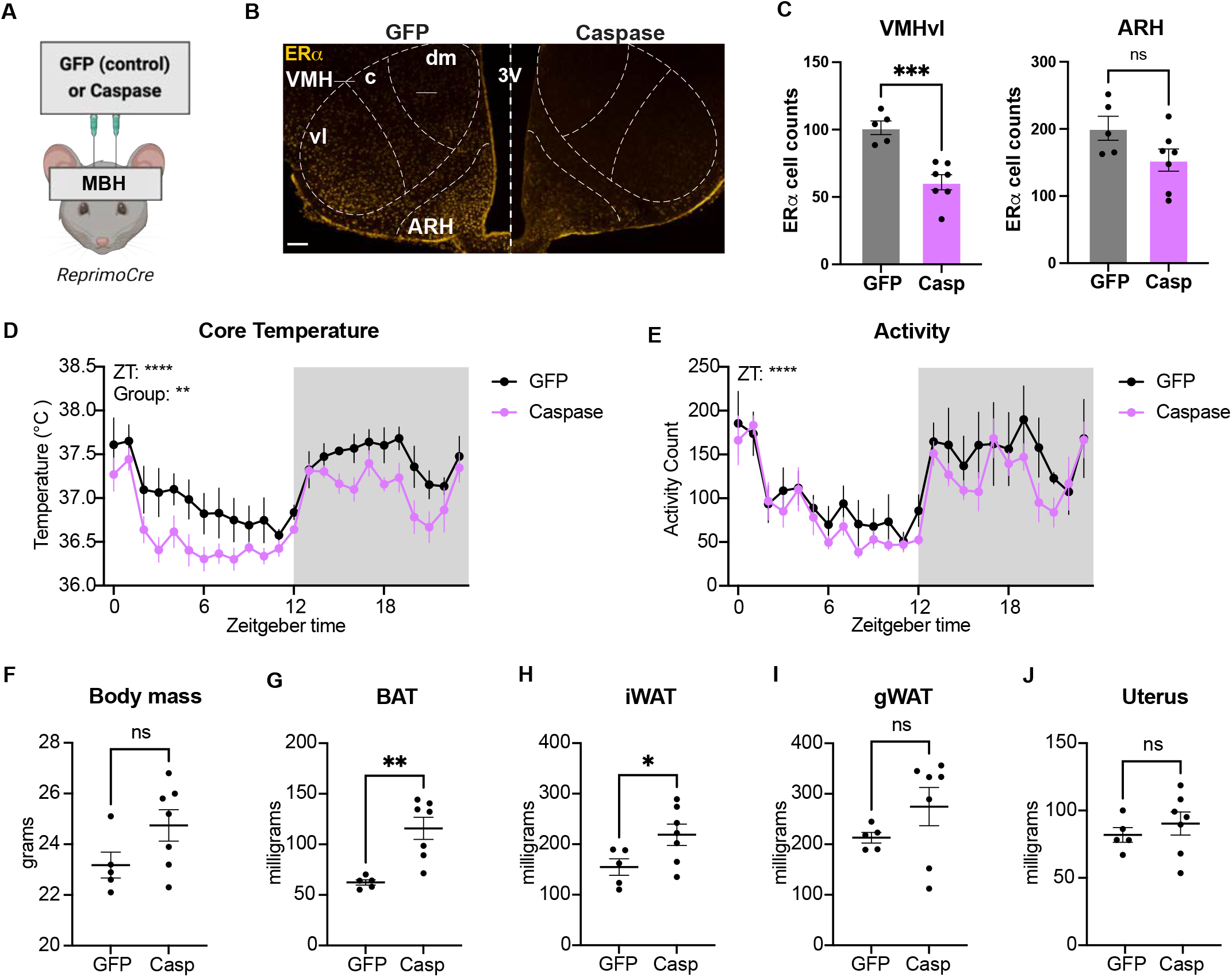
Ablating *Reprimo*+ cells in the MBH reduces core temperature in female mice. (A) Illustration depicting the stereotaxic delivery of a cre-dependent caspase or control virus targeting the VMHvl of female mice, with possible spread to other areas of the MBH. (B) Image of ERα immunoreactivity (yellow) in a control (GFP; left) and caspase-treated (casp; right) mouse. VMHvl delineated to aid in visualization. (C) Quantification of ERα+ cells in the VMHvl (left) and ARH (right) in control (gray bar) and caspase-treated (purple bar) female mice. (E, F) Core temperature and activity counts averaged hourly across 24 hours in control (black) and caspase-treated (purple) mice. (G) Body mass of control (GFP) and caspase-treated mice at 8 weeks of age. (H-K) Mass of dissected BAT (H), iWAT (I), gonadal WAT (J), and uterus (K) from GFP and caspase mice. N = 5 GFP and N = 7 caspase for all plots. Statistical analysis performed using mixed model Two-way ANOVAs (D-E) or t-tests (F-J). The mean ± SEM are depicted. Individual data points are shown in figures C-D and G-K.* *p* < 0.05, ** *p* < .01, **** *p* < 0.0001. Scale bar = 100 μm. VMH = ventromedial hypothalamus (c = caudal, dm = dorsomedial, and vl = ventrolateral areas), ARH = arcuate nucleus, 3V = third ventricle.

## Discussion

We investigated the role of *Esr1* in *Rprm*+ cells and the specific contribution of MBH^*Rprm*^ neurons on energy homeostasis. First, we characterized the *Rprm* lineage using a novel *Rprm*^*Cre*^ mouse and an *Ai14* reporter. We observed *Rprm*-driven tdTomato expression in the pancreas, uterus, ovary, pituitary, and brain. Interestingly, we did not observe evidence of *Rprm* expression in fat depots associated with heat generation or retention, including BAT, iWAT, or gWAT. It is intriguing that *Rprm* appears predominantly expressed in hormone-producing or hormone-transporting tissues (*i*.*e*., ovaries, pituitary, pancreas, and brain). In particular, these tissues are estrogen sensitive [47–49], raising the possibility that *Rprm* may play a role in the development or function of hormone-sensitive tissues. In contrast to the widespread expression of tdTomato, which labels the the *Rprm* lineage, we found that *Cre* expression in adults is more restricted and a reliable indicator of endogenous *Rprm* expression. The findings in adults validate the fidelity of the *Rprm*^*Cre*^ mouse line and suggest that tdTomato expression is a result of previous, transient *Rprm* expression in *Ai14*^*f/+*^;*Rprm*^*Cre*^ mice. Future studies will use tools that are spatially or temporally restricted, such as AAVs injected to specific tissues in adulthood, in *Rprm*^*Cre*^ mice to understand the role of cells or tissues that actively express *Rprm* at the time of manipulation.

Here we tested the role of *Esr1* in *Rprm*-lineage cells using *Rprm*^*Cre*^; *Esr1*^f/f^ (RERKO) mice. In the adult brain, *Esr1* and *Rprm* are co-expressed in at least four hypothalamic nuclei: the AVPV, the MPO, the VMHvl, and the ARH. RERKO mice had reduced ERα+ cell counts in all of these regions, except the MPO. These regions are associated with energy homeostasis and reproduction prompting us to compare these phenotypes in wildtype and RERKO mice.

As mice aged to four months, there was a subtle increase in body weight in males, but not females. However, there were no significant differences in fat mass, lean mass, or food consumption between RERKO mice and their WT siblings in male or female mice. We observed a reduction in ERα+ cells in the ARH but did not detect an effect on food intake, suggesting that the *Rprm* lineage does not substantially contribute to ERα cells in the ARH that modulate food intake. The most notable phenotype was a doubling of BAT mass that was specific to females.

This was surprising since previous studies on global *Esr1* KO mice report significant weight gain and adiposity, but no change in BAT mass [32,33]. Future studies could assess if there are differences in thermogenic capabilities of BAT tissue. In summary, while *Esr1* was knocked out in the *Rprm* lineage, which extends to multiple tissues, RERKO mice exhibited a more restricted phenotype than global *Esr1* KO.

Because we found the strongest effect of RERKO on the mass of thermogenic BAT, we compared the temperature of the core, BAT and tail skin. Interestingly, we found effects on core temperature only in females, not males, suggesting that estrogen signaling in the *Rprm* lineage is particularly important for maintaining normal temperatures in females. In females, there was no clear directionally of the temperature change; instead RERKO females appear to show reduced differences in temperature between day and night. A similar pattern is observed in BAT and tail temperature, although we only detected a main effect of genotype and no effect of the interaction between genotype and time. Thus, overall energy expenditure may be unchanged, as suggested by no change in body mass. Nonetheless, our results suggest that estrogen signaling in *Rprm* lineage cells may help maintain day-night patterns of core temperature by modulating both heat generation and dissipation processes.

In addition to differences in thermoregulatory patterns, we observed reproductive deficits in female RERKO mice: they did not cycle well and had abnormal reproductive anatomy. Thus, a limitation of this mouse model is potential differences in hormone levels, which may indirectly alter physiology and metabolism. To address this, we ovariectomized RERKO and WT mice and found persistent differences in core temperature, albeit with a different pattern than intact RERKO mice suggesting some hormonal effects.

Because we did not detect *Rprm* expression in BAT nor iWAT, we hypothesized that RERKO mice had altered central regulation of energy expenditure. Movement and BAT-mediated thermogenesis are associated with the VMHvl [10,11], making it the most promising mediator of a temperature phenotype. To conclusively test that *Rprm* cells in the brain, and not peripheral organs, regulate thermogenesis, we ablated *Rprm*+ cells in the MBH by stereotaxic delivery of a Cre-dependent caspase. We observed a ∼40% decrease in the number of ERα cells in the VMHvl, consistent with our prior observation and confirmation in this paper that 41% of ERα cells co-express *Rprm* [11]. Caspase-treated mice had lower core temperature than controls with no changes in movement, similar to previous effects of knocking down *Rprm* in the same region [11]. We also observed heavier BAT mass in caspase-ablated mice, suggesting an effect on BAT-mediated thermogenesis. Together, these studies suggest that MBH^*Rprm*^ cells regulate core body temperature. Taken together with the effects of ERα KO in the *Rprm* lineage, these findings suggest that MBH^*Rprm*^ cells may mediate some of the effects of estrogens on core temperature and thermogenesis. However, future studies will be required to definitively test this hypothesis using intersectional genetic approaches to specifically knock out *Rprm* in *Esr*1+ cells in the VMHvl or MBH.

## Conclusion

In summary, we use a new genetically engineered mouse model to show that estrogen-dependent regulation of cells in the *Rprm* lineage alters temperature, partly by modulating BAT thermogenesis and heat dissipating processes. Evidence of *Rprm* expression is not found in brown or white adipose tissue, suggesting that *Esr1* in *Rprm* cells within the brain (as opposed to in adipose tissue) is critical for proper thermoregulation. Specifically, it appears that *Rprm+* cells in the MBH drive patterns of heat generation via BAT. This work adds to a body of literature showing specialization of sub-populations of ERα neurons [10,13,22,24,50]. By identifying the cells driving regulation of thermogenesis, our research opens potential avenues for new precision treatments in metabolic disorders, such as obesity and menopause.

## Supporting information

Statistical tables

## Acknowledgements

The authors thank members of the Correa lab, particularly Paul Vander and Rachel Scott, for their feedback on the manuscript, as well as, Sakina Rashid and Hannah Azadi for technical assistance.

## Author contributions

JWP, LRC, and SMC conceptualized the present studies. JEV and LGK designed the *Rprm-Cre-FRT* mouse, and JWP designed the telemetry probe sleeves. JWP, LRC, NPS, ARV, AGB, HJF, MRH, EAD, WB, and KV contributed to experimental research and data collection. JWP, LRC, NPS, ARV, HJF, KV, and JIL provided data analysis and validation. SMC, LRC, and LGK provided funding and/or resources to complete the studies, and SMC provided mentoring and laboratory space. LRC and SMC wrote the manuscript with input from JWP, NPS, and JEV.

## Disclosures

### Data available

Data published in this article may be shared by emailing SMC.

