## Supplemental Figures for "Sex-specific thermoregulatory effects of estrogen signaling in *Reprimo* lineage cells"

**SUPPLEMENTARY MATERIALS**
**Supplementary Figures 1-6**
**Statistical Tables (separate excel file)**

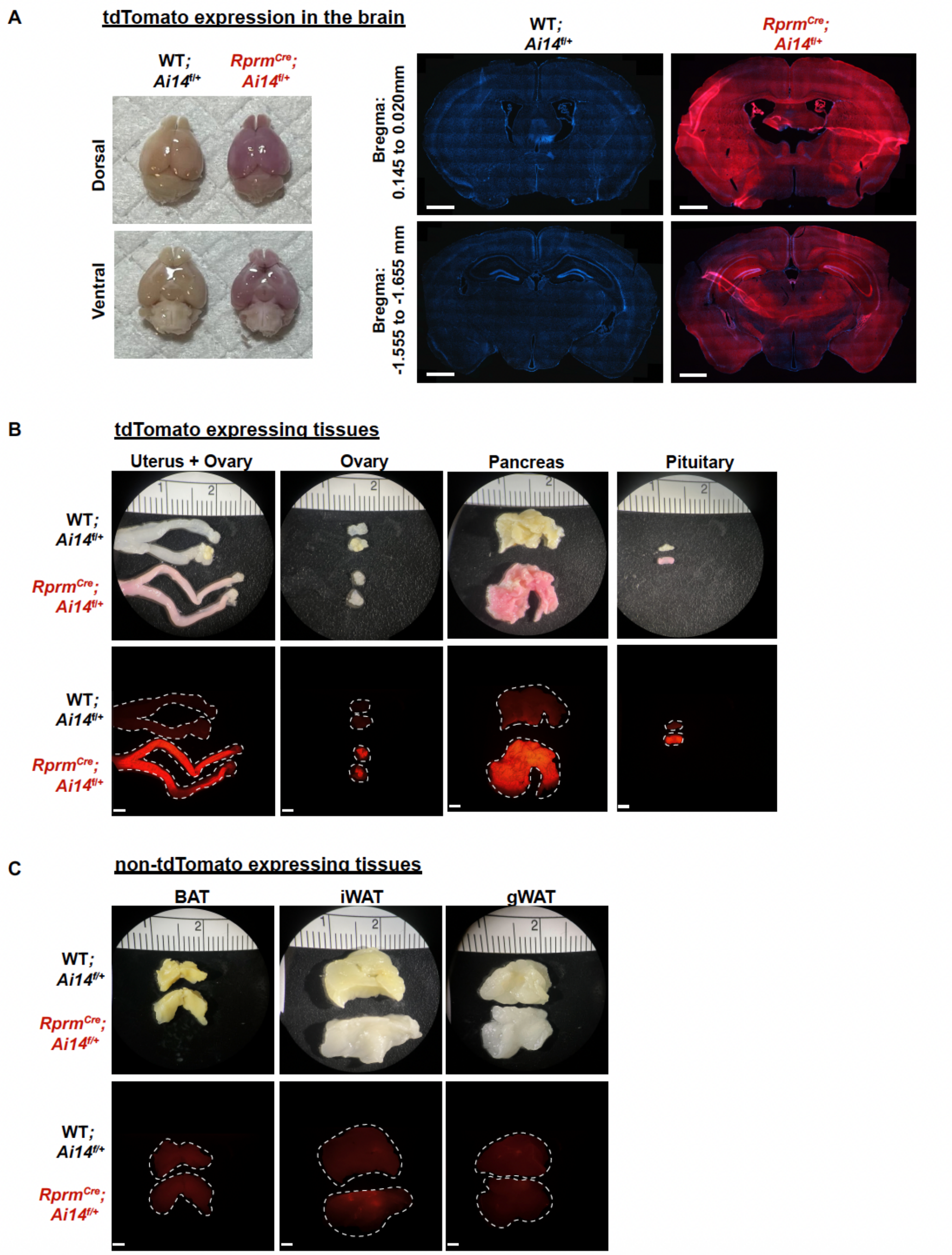


**Supplemental Figure 1: *Rprm* lineage tracing reveals robust expression in the brain and peripheral tissues.** Heterozygous $Rprm^{Cre}$ mice were bred with $Ai14^{f/f}$ mice to produce $Rprm^{Cre}$*;*$Ai14^{f/+}$ mice expressing tdTomato in *Rprm* lineage tissues. Cre-negative littermates (WT*;*$Ai14^{f/+}$ *mice*) were used to establish baseline tdTomato levels. (A-C) Representative images of tissues harvested from adult (13-15 weeks old) $Rprm^{Cre}$*;*$Ai14^{f/+}$ and WT*;*$Ai14^{f/+}$ mice. (A) Left – images highlighting tdTomato expression throughout the whole brain in male mice. Right – 30 µm DAPI-stained coronal brain sections emphasize the expression of tdTomato in the cortex, thalamus, and hypothalamus across representative bregma levels. Scale bars = 1 mm. (B) Peripheral tissues exhibiting tdTomato expression include the uterus, ovary, pancreas, and pituitary in female mice. (C) Peripheral tissues that do not express tdTomato include adipose tissue depots (BAT, iWAT, and gWAT) in female mice. Brightfield images displayed on top, fluorescent images below. Scale bars = 200 µm. BAT = brown adipose tissue, iWAT = inguinal white adipose tissue, gWAT = gonadal white adipose tissue.

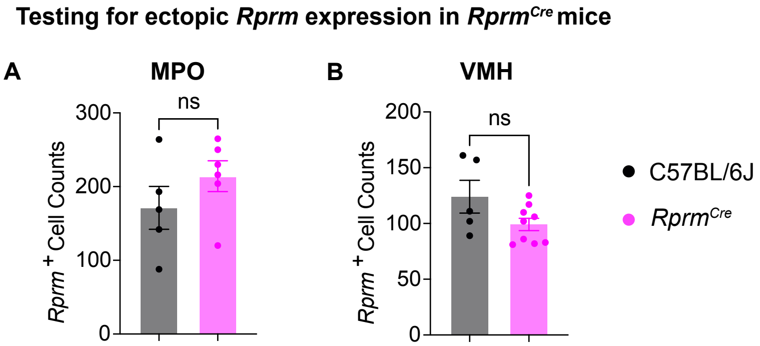


**Supplemental Figure 2: Knock-in construct does not alter *Rprm* expression in the MPO and VMH.** *Rprm*+ cell counts were compared between *Rprm*$^{Cre}$ and wildtype C57BL/6J mice (n=5-9 per group) in the MPO (A) and VMH (B). Statistical analysis performed using Welch's t-test.

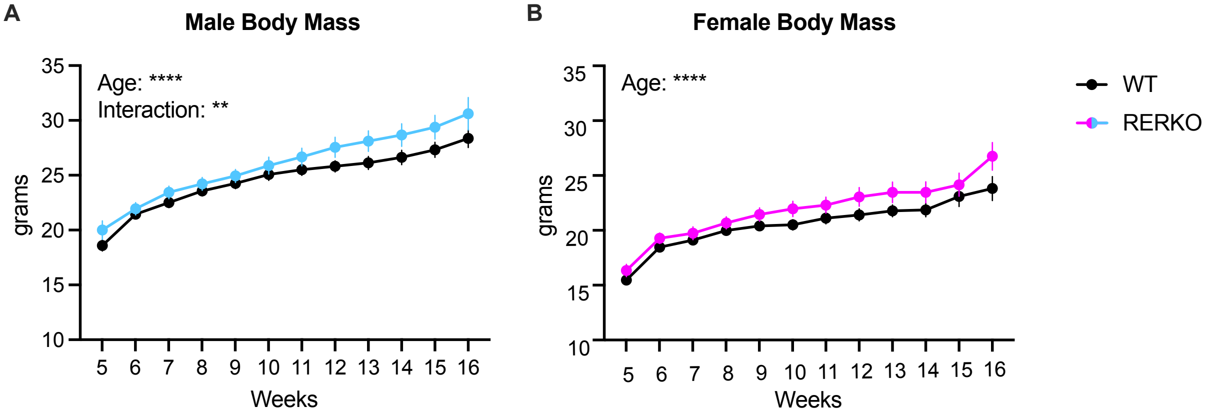


**Supplementary Figure 3. Longitudinal body weight analysis of RERKO mice.** (A) Body mass measurements of control (black) and RERKO (blue) male mice from 5 to 16 weeks of age (n = 16 WT, n = 18 RERKO). (B) Body mass measurements of control (black) and RERKO (pink) female mice from 5 to 16 weeks of age (n = 13 WT, n = 13 RERKO). In females, we only detected an effect of age ($F_{11,238}$ = 61, $p < 0.0001$). Mean ± SEM is plotted at each time point. A mixed-effects model using Restricted Maximum Likelihood (REML) estimation was used for statistical analysis due to missing values.

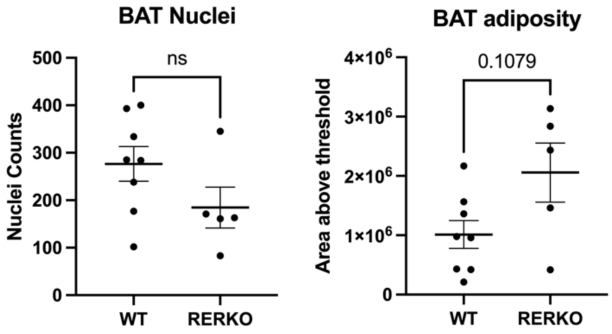


**Supplementary Figure 4. BAT nuclei and BAT adipose quantification in control and RERKO female mice.** A) Nuclei within a fixed area of BAT tissue were counted in 40x images in control and RERKO adult female mice. B) To roughly compare the adiposity in BAT tissue, the total area of the white portion of BAT images (a proxy for fat droplets) was also estimated using thresholding analysis in ImageJ. Statistical analysis performed using t-test.

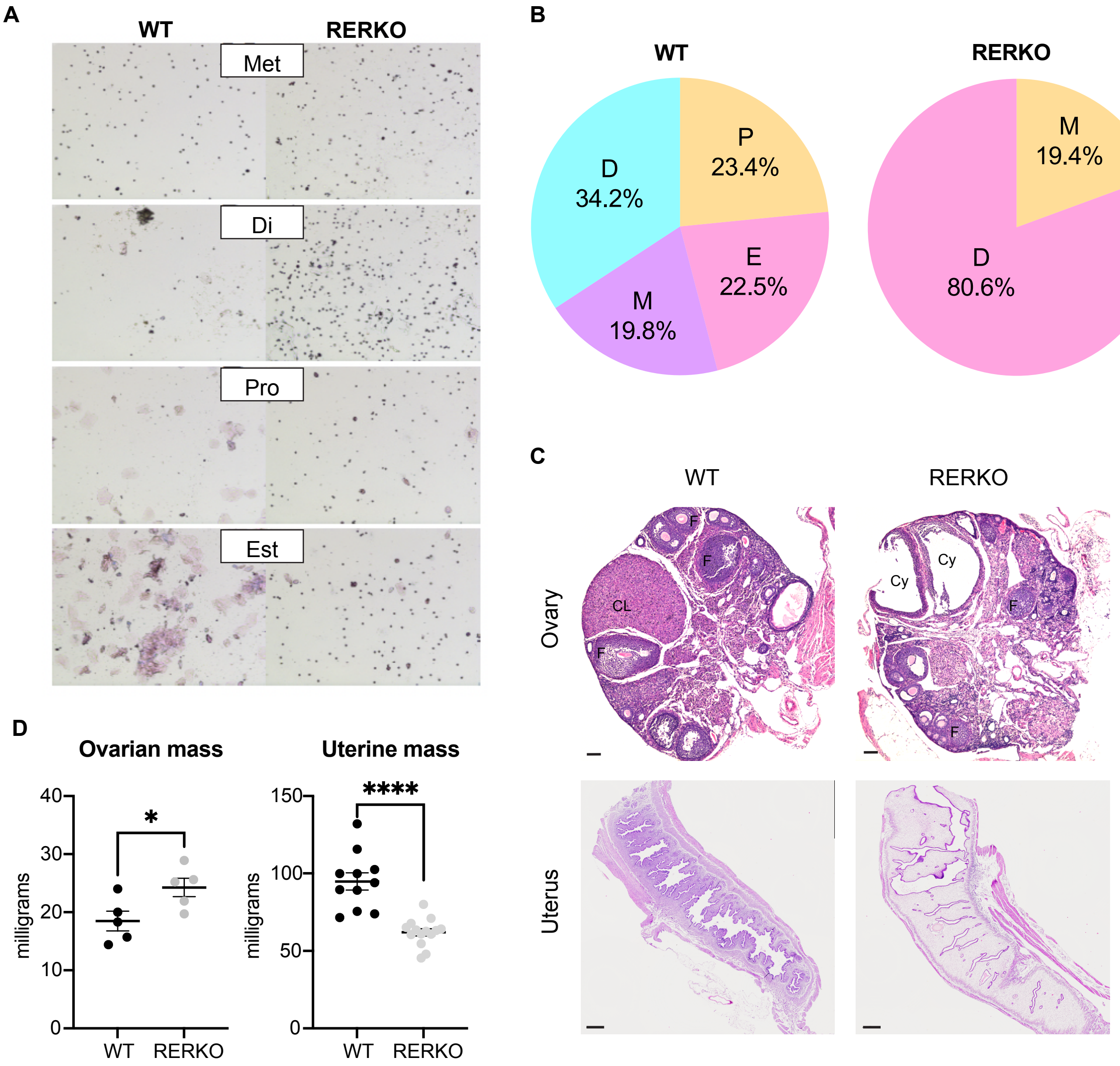


**Supplementary Figure 5. RERKO female mice have disrupted reproduction.** (A) Images depict vaginal lavages from WT (left) and RERKO (right) mice stained with Giemsa. While the WT mice display expected cellular morphological changes across the estrous cycle, swabs from RERKO mice mostly contain leukocytes. (B) The percent of days across 14 days spent in diestrus [D], proestrus [P], estrus [E], and metestrus [M] in WT (left) and RERKO (right) mice. (C) Ovarian histology visualized with hematoxylin & eosin stain. Follicular cysts are present and the corpus luteum is absent in the ovary from RERKO mice. CL = corpus luteum, F = ovarian follicles, Cy = cyst. Scales are 100 µm (ovary) and 500 µm (uterus). D) Dissected ovarian and uterine mass from control (black) and RERKO (gray) mice. T-test used to compare groups. The mean ± SEM are depicted. Individual data points are shown in Figure D. * $p < 0.05$, **** $p < 0.0001$.

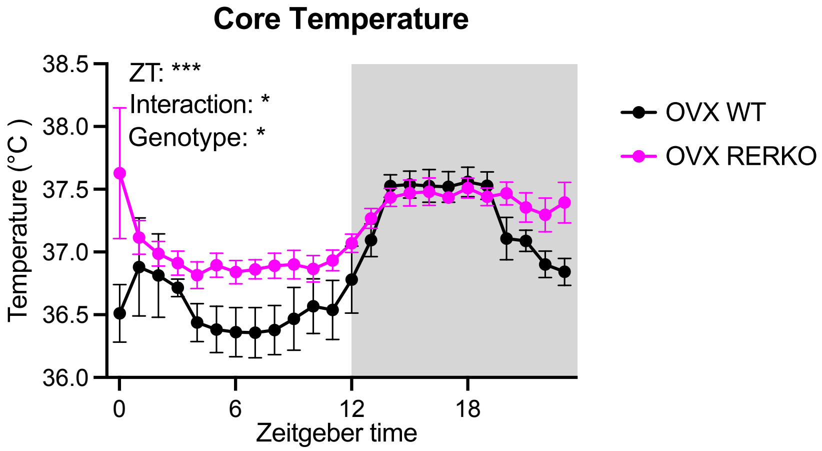


**Supplementary Figure 6. Core temperature in RERKO mice lacking ovaries.** Core temperature averaged hourly across 24 hours in ovariectomized (OVX) WT littermates (black) or RERKO (pink) mice. Two-way mixed ANOVAs used for statistical analysis. The mean ± SEM are depicted for each hour. * $p < 0.05$. ZT = Zeitgeber time. C = Celsius. Shaded box indicates the dark (active) phase of the day.
